# EXOSOMES FROM CYCLIC MICE MODULATE LIVER TRANSCRIPTOME IN ESTROUPAUSE MICE INDEPENDENT OF AGE

**DOI:** 10.1101/2024.11.04.621842

**Authors:** Bianka M. Zanini, Bianca M. Ávila, Jéssica D. Hense, Driele N. Garcia, Sarah Ashiqueali, Pâmela I. C. Alves, Thais L. Oliveira, Tiago V. Collares, Miguel A. Brieño-Enríquez, Jeffrey B. Mason, Michal M. Masternak, Augusto Schneider

## Abstract

**Background:** Exosomes are extracellular vesicles secreted by cells that contain microRNAs (miRNAs). These miRNAs can induce changes in gene expression and function of recipient cells. In different cells exosome content can change with age and physiological state affecting tissues function and health.

**Aims:** Therefore, the aim of this study was to characterize the miRNA content and role of exosomes from cyclic female mice in the modulation of liver transcriptome in estropausal mice.

**Main Methods:** Two-month-old female mice were induced to estropause using 4-vinylcyclohexene diepoxide (VCD). At six months of age VCD-treated mice were divided in control group (VCD) and exosome treated group (VCD+EXO), which received 10 injections at 3-day intervals of exosomes extracted from serum of cyclic female mice (CTL).

**Key findings:** Exosome injection in estropausal mice had no effect on body mass, insulin sensitivity or organ weight. We observed ten miRNAs differentially regulated in serum exosomes of VCD compared to CTL mice. In the liver we observed 931 genes differentially expressed in VCD+EXO compared to VCD mice. Interestingly, eight pathways were up-regulated in liver by VCD treatment and down-regulated by exosome treatment, indicating that exosomes from cyclic mice can reverse changes promoted by estropause in liver. *Cyp4a12a* expression which is male-specific was increased in VCD females and not reversed by exosome treatment.

**Significance:** Our findings indicate that miRNAs content in exosomes is regulated by estropause in mice independent of age. Additionally, treatment of estropausal mice with exosomes from cyclic mice can partially reverse changes in liver transcriptome.

## Introduction

Exosomes are nanometric extracellular vesicles that can be found in the bloodstream and transfer molecular signals from tissue to tissue [1-3]. Exosomes can carry active biological content, including proteins, lipids, and RNAs [4,5], which is influenced by their origin and the physiological condition of the secretory cell [6,7]. Small RNAs inside exosomes can induce changes in the activity of recipient cells, suggesting an endocrine role [8]. Exosomes isolated from serum of young mice can modulate expression of age-related genes when injected into old mice [9-11]. Therefore, it is suggested that exosomes play an active role in the regulation of aging. In the heterochronic parabiosis model, the circulatory systems of young and old mice are joined, resulting in slower aging in the older mice, as indicated by transcriptome and epigenomic changes [12]. Similarly, treatment of older rats with plasma from younger rats prevented age-related epigenetic changes [13]. Overall, this evidence suggests that blood derived factors can affect age-related pathways and therefore, the role of exosomes and its contents should be better explored.

MicroRNAs (miRNAs) are small RNAs (19-25 nucleotides) that regulate gene expression by modulating the stability and translation of messenger RNA (mRNA) [14]. The content of miRNAs in blood cells [15], serum [16] and blood derived exosomes [17] changes with age in mice. Therefore, exosomal miRNAs (exo-miRs) have been proposed as non-invasive biomarkers of aging and age-related diseases [18,19]. In addition, exo-miRs can have an active role in different physiological conditions due to their potential to modulate gene expression in target cells [20]. In this sense, during female aging there is also a decline in ovarian activity that leads to hormonal changes and infertility. The ovarian reserve of oocytes is finite and its exhaustion results in the end of reproductive life, known as menopause in women [21] and estropause in mice [22]. Early onset of menopause is associated with accelerated epigenetic age [23] and increased mortality from various causes [24]. This suggests that the ovary can actively influence overall female health. In mice, ovariectomy reduces lifespan [25], while the transplantation of young ovaries into old females increases it [26]. Interestingly, the transplantation of young ovaries with depleted germ cells further increased lifespan of recipients [27], indicating that factors other than hormones produced by young ovaries may be responsible for this protective effect. We showed before that chemically induced estropause in young mice changed the profile of serum exo-miRs [28]. Similarly, post-menopausal women have a different profile of serum exo-miRs compared to pre-menopausal women [29]. This suggests that changes in serum exo-miR profile may regulate metabolism in females experiencing estropause and can be targets for interventional studies.

The evidence presented suggests that the young ovary produces intrinsic factors that can prevent aging, which becomes especially important when ovarian activity is reduced. Given that most women undergo a natural transition to menopause while retaining ovarian tissue, the compound 4-vinylcyclohexene diepoxide (VCD) can induce follicular depletion with retention of ovarian tissue in mice, simulating the changes observed during menopausal transition in women [30,31]. VCD mice have increased body weight gain, although no effect on serum lipid and liver damage markers were observed [32,33]. Nevertheless, VCD induced estropause changes the hepatic transcriptome, affecting lipid biosynthesis related pathways [33], and increaseing the susceptibility to liver damage under a high-fat high-sucrose diet [33]. We also observed increased liver reactive oxygen species under regular chow diet in VCD treated estropausal mice compared to cyclic mice [32]. Changes in lipid synthesis are critical, as post-menopausal women have higher serum cholesterol levels [34] and a greater risk of cardiovascular diseases [35]. Therefore, it is important to understand the regulation of liver transcriptome in response to interventions in estropausal mice. The use of VCD mice also allows us to measure the effects of ovarian failure independent of age, as the effects of naturally occurring estropausal mice are confounded with that of age itself.

Exosomes can have therapeutic potential due to their low immunogenicity and toxicity and serve as a drug delivery platform [36]. Advancements in nanotechnology enable the encapsulation of therapeutic agents in exosomes, such as chemotherapeutic drugs, small molecules, miRNAs, and siRNAs [36,37]. Although there are still many challenges for its clinical use, such as large-scale manufacturing, cell sources, and target specificity, exosome engineering remains a promising therapeutic strategy for treatment of several diseases. The administration of exo-miRs also has been shown to have therapeutic potential in neurodegenerative [38], inflammatory [39] and cardiovascular diseases [40]. Therefore, our aim was to characterize the miRNA content, and the role of exosomes derived from cyclic females in modulating liver transcriptome in estropausal mice independent of age.

## Methodology

### Animals and experimental design

This study was approved by the Animal Experimentation Ethics Committee from Universidade Federal de Pelotas (number 1379560). For this study C57BL/6 female mice (2-month-old, n=101) were kept under controlled conditions of temperature and light (22± 2°C, cycles 12 hours light/12 hours dark) and provided with *ad libitum* access to a standard chow and water. A subset of mice (n=21) was treated daily with intraperitoneal injection of VCD (160 mg/kg) for 20 days to induce estropause [41,42], while a subset of control females (n=10) received placebo (sesame oil) at the same frequency. At six months of age, estropause and cyclic (CTL) mice were randomly allocated to a control and experimental group to receive exosomes (VCD+EXO) or placebo (VCD) injection **(Figure 1).** After 48 h of the last exosome injection, mice were anesthetized using isoflurane and euthanasia was performed following a 4-hour fasting. Exsanguination was performed by cardiac puncture followed by cervical dislocation. Mice were dissected, and ovaries, white adipose tissue depots and liver were collected and stored in 10% buffered formalin and at -80oC. Blood samples werecentrifuged at 5,000xg for 10 min for serum separation. The amount of fat tissue was determined by weighing the amount dissected relative to body mass immediately after euthanasia.

**Figure 1.**
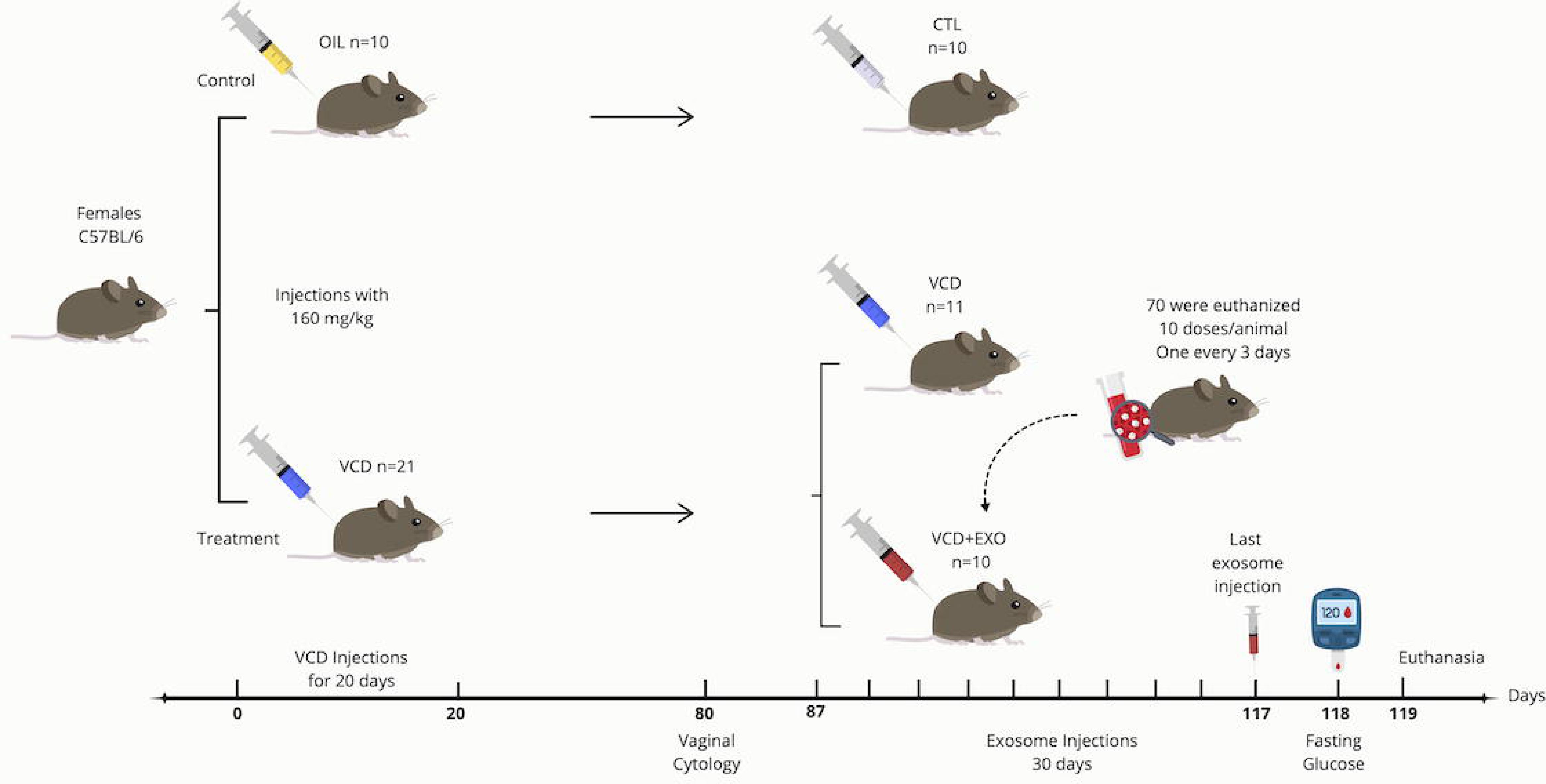
Experimental Design. Female mice at 60 days of age received injections of sesame oil or 4-vinylcyclohexene diepoxide (VCD) for 20 consecutive days. Sixty days after the end of the VCD/oil injection, estrous cycle assessment was conducted to confirm estropause. After the VCD+EXO group started exosome therapy every three days via intraperitoneal injection for 30 days. One day before the end of therapy, all groups underwent an insulin tolerance test, and in the next day, euthanized.

### Exosome isolation and injection

At six months of age, untreated cyclic control females C57BL/6 mice (n=70) were euthanized to collect serum for exosomes extraction. Exosomes were isolated from serum using a commercial kit (Total Exosome Isolation Reagent from serum, Invitrogen, Vilnius, Lithuania) following the manufacturer’s protocol. Briefly, a buffer was added to the serum samples and incubated at 5°C for 30 minutes. Precipitated exosomes were recovered by centrifugation at 10,000 xg for 10 minutes. The pellet was resuspended in phosphate-buffered saline (PBS).

Total protein was extracted from exosomes and measured using a commercial kit (Pierce BCA Protein Assay, Thermo Scientific, Carlsbad, CA, USA) according to the manufacturer’s instructions. Exosomes extracted from the serum of mice resulted in approximately 1000 μg of protein. Therefore, we standardized the injection concentration to 800 μg/mouse, similar to a previous study using exosomes isolated from liver [43]. Each treated mouse therefore received 80% of serum exosome content from one donor mouse at each injection timepoint. Exosomes extracted from serum of individual cyclic mice were combined into a single tube for dilution. Exosomes aliquots were then prepared and stored in -80°C until the moment of injection. Each mouse in the VCD exosome treated group received one injection every three days, in a total of 10 injections/mice in a 30-day interval (8 ug/μL; 100 μL/mouse).

### Exosome Characterization

Particle concentration and size distribution of extracted exosomes from cyclic mice were determined by nanoparticle tracking analyzer using a NanoSight NS300 with 488 nm laser set at 25°C and camera level 11 (Malvern instruments). The NanoSight sample pump was used in conjunction with 1 mL syringes at flow speed of 30. Samples were run immediately after dilution in 0.1uM filtered HPLC grade water. Each analysis consists of five 60-second video captures per sample (University of Florida – ICBR, Cytometry Core Facility; RRID:SCR_019119).

Part of the isolated exosomes was stained with SYTO™ RNASelect™ Green Fluorescent Cell Stain (Invitrogen), following the manufacturer’s instructions. Exosomes were stained one day before administration as overnight incubation was necessary. Mice (2 VCD+EXO and 1 CTL) were euthanized 24 hours post-injection and blood was collected freshly fixed on slides and subsequently visualized using a confocal fluorescence microscope (FluoView 1000; Olympus).

### Vaginal Cytology

Vaginal smears were obtained 60 days after the end of VCD injections in CTL, VCD and VCD+EXO groups (n=31). Daily smears were performed at the same time for five consecutive days. The collected material was stained using rapid panoptic. Acyclic females for five days were considered to be in estropause.

### Insulin Tolerance Test

An insulin tolerance test (ITT) was performed one day before euthanasia. For this, mice received i.p. injection of insulin (0.5 IU/kg body weight) after a 2-hour fast. Blood was collected 0, 5, 20, and 35 minutes after injection, and glucose levels were measured using a glucometer. The percentage of glucose decay between 5 and 20 minutes after insulin injection was calculated.

### Follicle counting

The ovaries were removed from 10% buffered formalin, dehydrated in alcohol, cleared in xylene, embedded in paraffin, and then cut into 5 μm sections using a semiautomatic microtome (RM2245 38, Leica Biosystems Newcastle Ltd., Newcastle Upon Tyne, UK). One out of every six sections were selected and placed on standard histological slides. After drying in a 56°C oven for 24 h, the slides were stained with hematoxylin and eosin and mounted with coverslips and synthetic resin (Sigma Chemical Company®, St. Louis, MO, USA). Images of the ovarian sections were captured using a microscope (Nikon Eclipse E200, Nikon Corporation, Japan) with 4×, 10×, and 40× objectives. Follicles with clearly visible oocyte nuclei were quantified. The number of follicles counted was divided by the total number of sections for each ovary. The follicle classification protocol was based on previous protocol [44].

### miRNA Library Preparation, Sequencing and Processing

For exosome RNA extraction, a commercial kit for total RNA and protein isolation from exosomes (Total Exosome RNA & Protein Isolation Kit, Invitrogen) was used. Total RNA was measured using the Qubit equipment (ThermoFisher).

miRNA libraries were prepared from total RNA extracted from serum exosomes of CTL, VCD and VCD+EXO mice (n=3/group) using the NEXTFLEX® Small RNA-Seq Kit V4 (Perkin Elmer, Seer Green, Beaconsfield, United Kingdom) and submitted to sequencing (Illumina NovaSeq X Plus). Alignment and quantification of miRNA libraries was performed using cutadapt and bowtie2 (Martin 2011) as described before [45]. Statistical analyses for differentially expressed miRNAs were performed using the software R (4.3.2) and the Bioconductor package EdgeR. Read counts were normalized for library depth, and pairwise comparisons, measuring fold change, uncorrected P-values from the negative binomial distribution, and adjusted P values (false discovery rate; FDR) were obtained. miRNAs with an FDR<0.1 and fold change (FC) >2.0 were considered upregulated; and with FDR<0.1 and FC<0.5 were considered down-regulated. Principal components analysis (PCA) and heatmap analysis were also performed to observe sample distribution using SRPlot [46]. Raw miRNA sequencing data is available from NCBI SRA PRJNA1171446.

The mirPath tool (version 3.0) was used to predict target genes of the differentially regulated miRNAs using the microT-CDS v. 5.0 database [47] and for retrieving KEGG molecular pathways [48,49], considering P<0.05 as significant. For pathway analysis, miRNAs were analyzed separately as down-or up-regulated.

### mRNA Library Preparation, Sequencing and Processing

Liver tissue was collected from CTL, VCD and VCD+EXO (n=3/group) mice 48 h after the last exosome injection, and total RNA was extracted using the TRIzol protocol. RNA was dissolved in RNAse free water, and its concentration and quality were estimated by spectrophotometry (EpochTM Microplate Spectrophotometer, BioTek, Winooski, VT, USA). Libraries were prepared using poly-A selected mRNA library using the xGen RNA library preparation kit (Integrated DNA Technologies, Coralville, IA, USA) with NEBNext® Poly(A) mRNA Magnetic Isolation Module (New England Biolabs, Ipswich, MA, USA). Library sequencing was also performed in a NovaSeq X Plus equipment.

The mapping of sequencing reads to the mouse transcriptome was performed using HiSat2 (Pertea, Kim et al. 2016). The number of reads aligned to its corresponding gene was calculated by HTSeq 0.6.1 [50]. Statistical analyses for differentially expressed mRNAs were performed using the software R (4.3.2) and the Bioconductor package EdgeR using the HTSeq output count. Read counts were normalized for library depth, and pairwise comparisons, measuring fold change, uncorrected P-values from the negative binomial distribution, and FDR were obtained. Genes with FDR<0.1 and FC>2.0 were considered upregulated; and with FDR<0.1 and FC<0.5 were considered down-regulated. Principal components analysis (PCA) and heatmap analysis were also performed to observe sample distribution using SRPlot [46]. mRNAs were further processed for pathway analysis using the Generally Applicable Gene-set Enrichment (GAGE), which uses log-based fold changes as per gene statistics, and Pathview packages in R (Luo and Brouwer 2013). P<0.05 were considered significant for pathways and GO Terms analysis. Raw mRNA sequencing data is available from NCBI SRA PRJNA1171446.

### Statistical Analyses

Statistical analyses were performed using GraphPad Prism 10. One-way ANOVA was used for body/organ mass, follicle count, and glucose metabolism. Repeated measures ANOVA was used for glucose levels during ITT analysis. A P<0.05 was considered statistically significant.

## Results

### Estropause confirmation

All females injected with VCD were acyclic before the start of exosome treatment, confirmed by vaginal cytology. The number of follicles in all stages of development was reduced in estropause mice **(Suppl. Fig. 1)**. No secondary and tertiary follicles were observed in estropause females **(Suppl. Fig. 1**), confirming all VCD injected females were acyclic. Exosomes had no effect on follicle numbers.

### Body composition and insulin metabolism

Mice in estropause had higher body mass compared to cyclic mice before the start of exosomes treatment (**Fig. 2A**; p=0.04). Exosomes treatment did not affect body mass change during treatment (**Fig. 2B**). The estropause groups (VCD and VCD+EXO) had lower uterine mass (**Fig. 2C**; p<0.0001), as well as lower peri-gonadal WAT (**Fig. 2D**; p=0.01). Visceral WAT (**Fig. 2E**; p=0.07), peri-renal WAT (**Fig. 2F**; p=0.08), kidney (**Fig. 2G**; p=0.65), and liver mass (**Fig. 2H**; p=0.18) were not different among groups.

**Figure 2.**
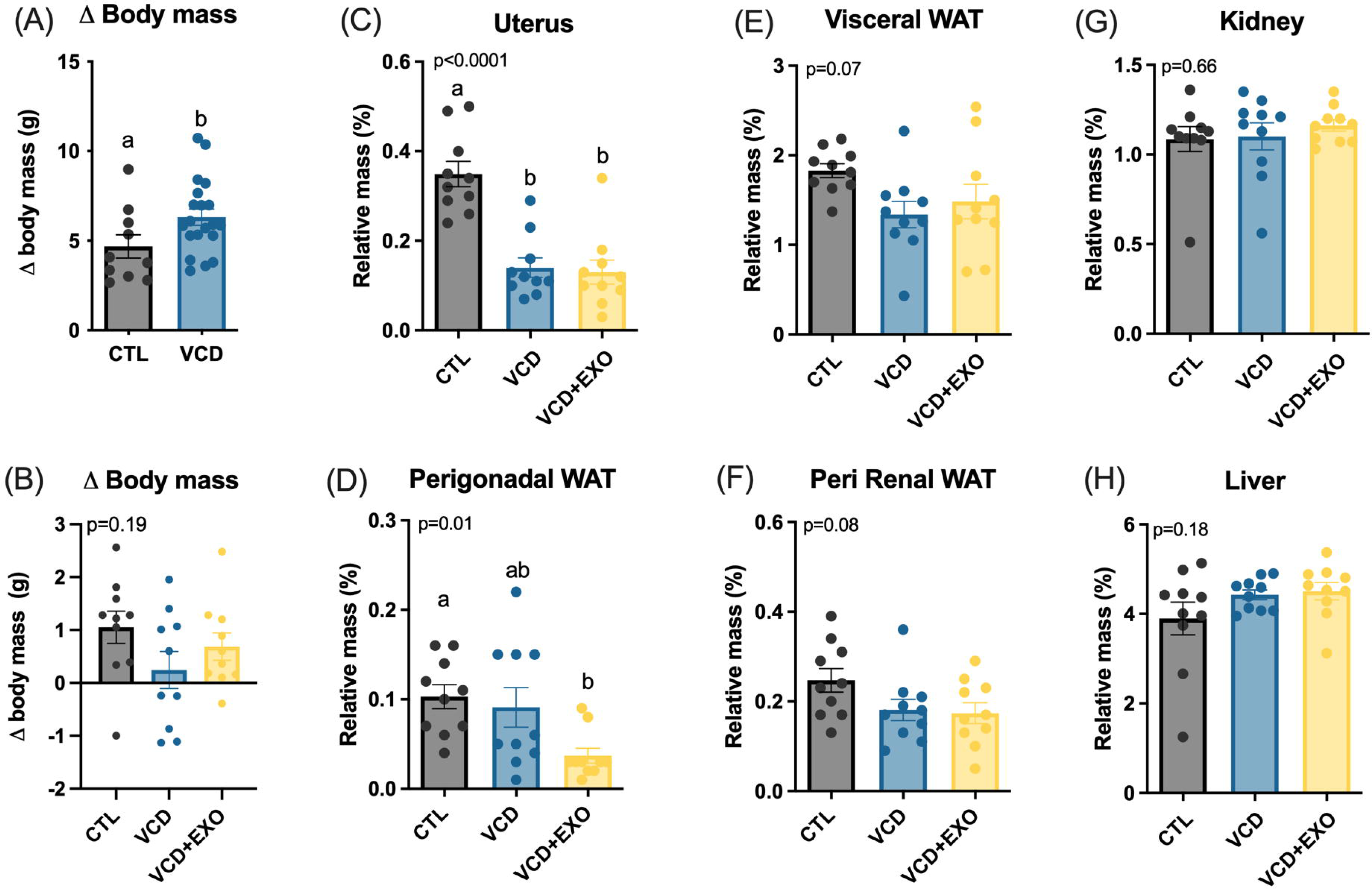
Body and organ mass. (A) Body mass change from the end of VCD treatment until the beginning of exosome treatment (B) Body mass change during the 30 day period of exosome injection. Relative mass of (C) uterus (D) perigonadal white adipose tissue (WAT); (E) visceral WAT; (F) perirenal WAT; (G) kidney; and (H) liver for control (CTL), estropause placebo (VCD) and estropause exosome (VCD+EXO) mice. Values are represented as mean ± standard error of the mean (SEM). Different letters indicate statistical difference at p<0.05).

Higher basal glucose levels were observed in the VCD+EXO (**Fig. 3A**; p=0.02) compared to CTL group. However, during the ITT, the rate of glucose decay was not different between groups (**Fig. 3B and C**; p=0.38).

**Figure 3.**
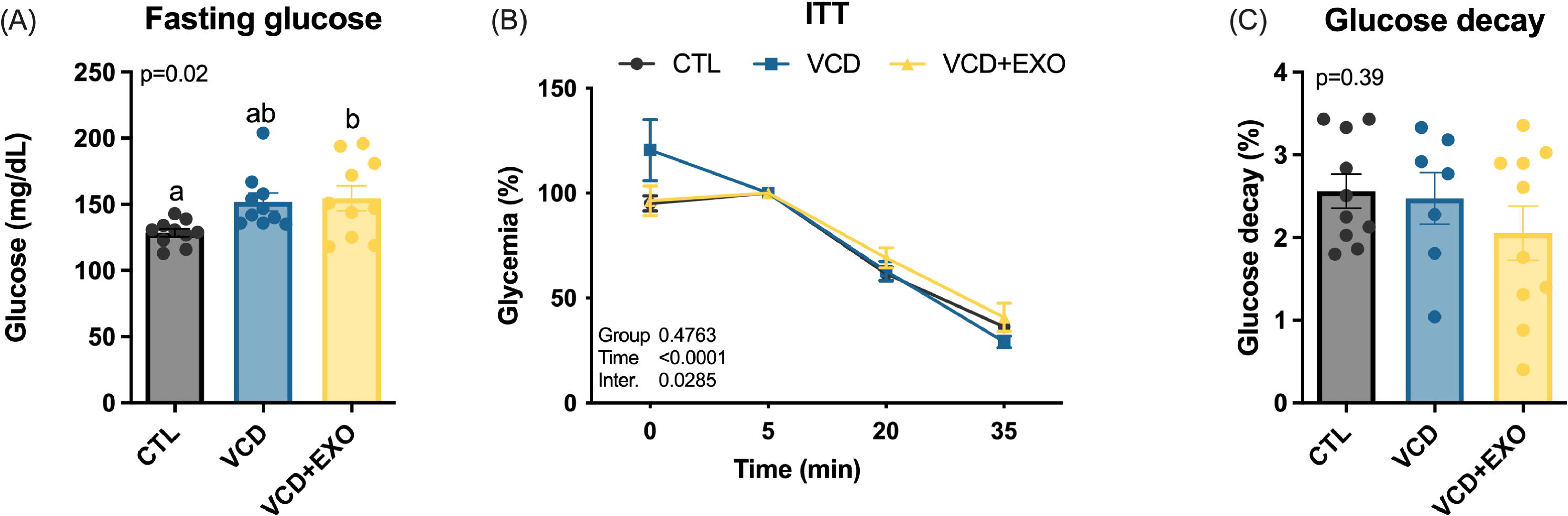
Glucose metabolism. (A) Fasting glucose levels for control (CTL), estropause (VCD) and estropause receiving exosome (VCD+EXO) mice. (B) Insulin tolerance test (ITT) and(C) glucose decay constant (KITT). Values are represented as mean L standard error of the mean (SEM). Values are represented as mean ± standard error of the mean (SEM). Different letters indicate statistical difference at p<0.05).

### Exosome characterization

We observed isolated and stained exosomes in blood smears 24 hs after i.p. injection by fluorescent emission **(Fig. 4A-B)**. The analysis of exosome particles was also performed in isolated exosomes before injection. Exosomes had an average diameter of 166.4 ± 2.3 nm, and the concentration was 1.68^10^ ± 9.31^8^ particles/mL **(Fig. 4C-E)**. The protein content of pooled exosomes was 1178 µg/mL and 78 ng/ml of RNA.

**Figure 4.**
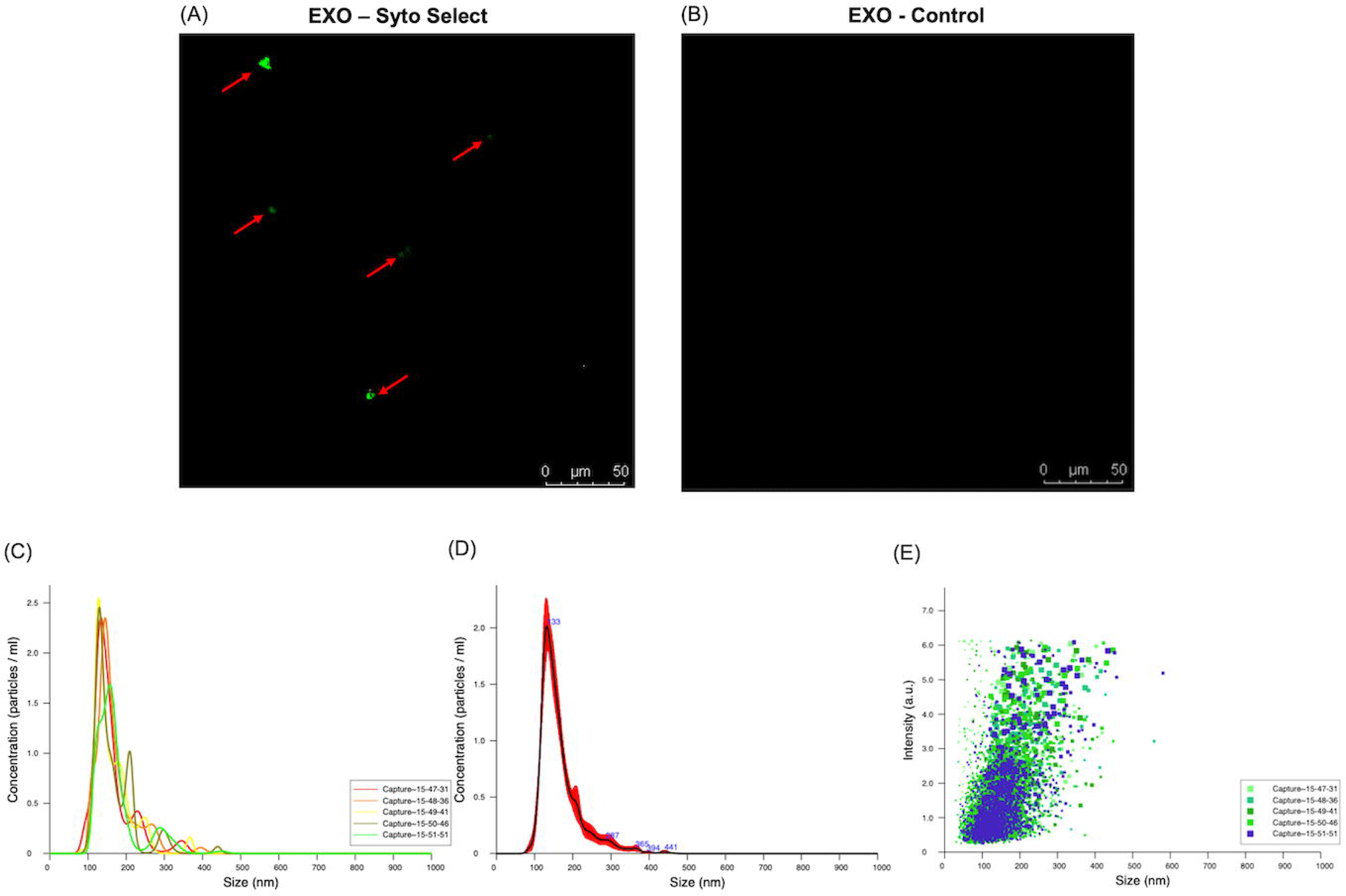
Exosome characterization. (A) Image of stained exosomes in blood smears 24 h post-injection in treated mice; (B) control blood smear without exosome staining; Exosome nanoparticle tracking analysis using Nano Sight NS300 system indicating particle size and concentration (A and B) and intensity (C).

### miRNA Serum Sequencing

A total of 355 miRNAs were detected in serum exosomes. We could observe that CTL samples clustered distinctively from VCD and VCD+EXO samples in the PCA analysis **(Fig. 5A)**. In the heatmap analysis we could also observe that VCD and VCD+EXO samples were mixed but distinct from CTL samples **(Fig. 5B)**. The number of differentially regulated miRNAs among CTL, VCD, and VCD+EXO groups is shown in the Volcano plots and Venn diagram **(Fig. 6).** We could not observe any difference between VCD and VCD+EXO mice in the serum miRNA profile, indicating that exosome treatment was not effective in sustained changes in the serum miRNA profile 48 hs after injection. The full list of regulated miRNAs is presented in **Suppl. Table 1**. miR-103-3p (down), miR-320-3p (down), miR-20a-3p (up), miR-150-5p (up), miR-181a-1-3p (up), miR-297a-3p (up) and miR-345-5p (up), miR-466f-3p (up) were commonly regulated by estropause and not reversed by the exosome treatment. Next, we examined the pathways regulated by the predicted target genes of regulated miRNAs among groups. We observed 46 pathways for down-regulated miRNAs in VCD+EXO compared to the CTL group, while VCD group had only 11 pathways down-regulated compared to the CTL group (**Suppl. Table 2**).

**Figure 5.**
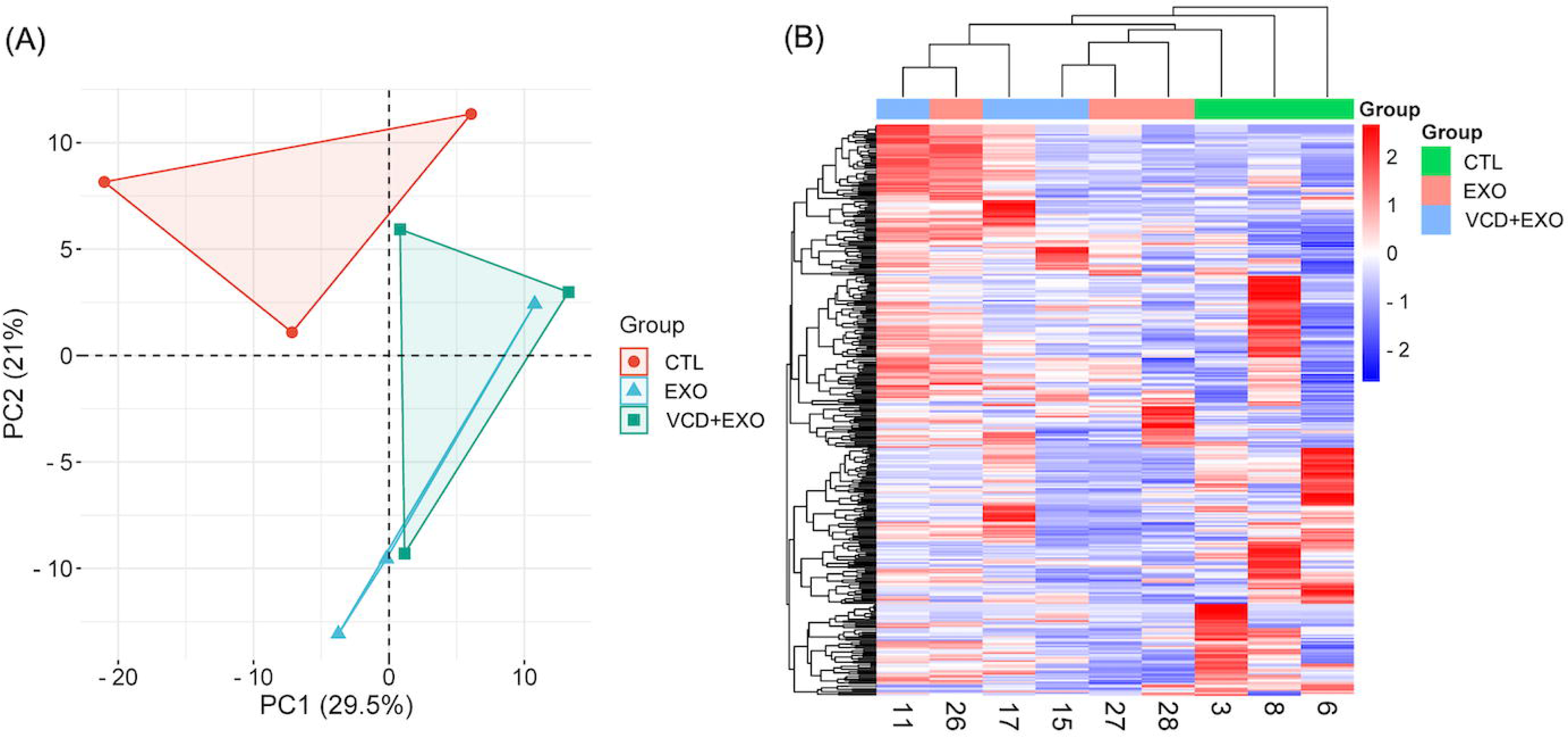
Serum exosomal microRNA (miRNA) analysis. (A) Principal component analysis of serum exosomal miRNAs for control (CTL), estropause (VCD) and estropause exosomes (VCD+EXO) mice; (B) Unsupervised hierarchical clustering ordered by the adjusted level of miRNA expression for all expressed miRNA in serum.

**Figure 6.**
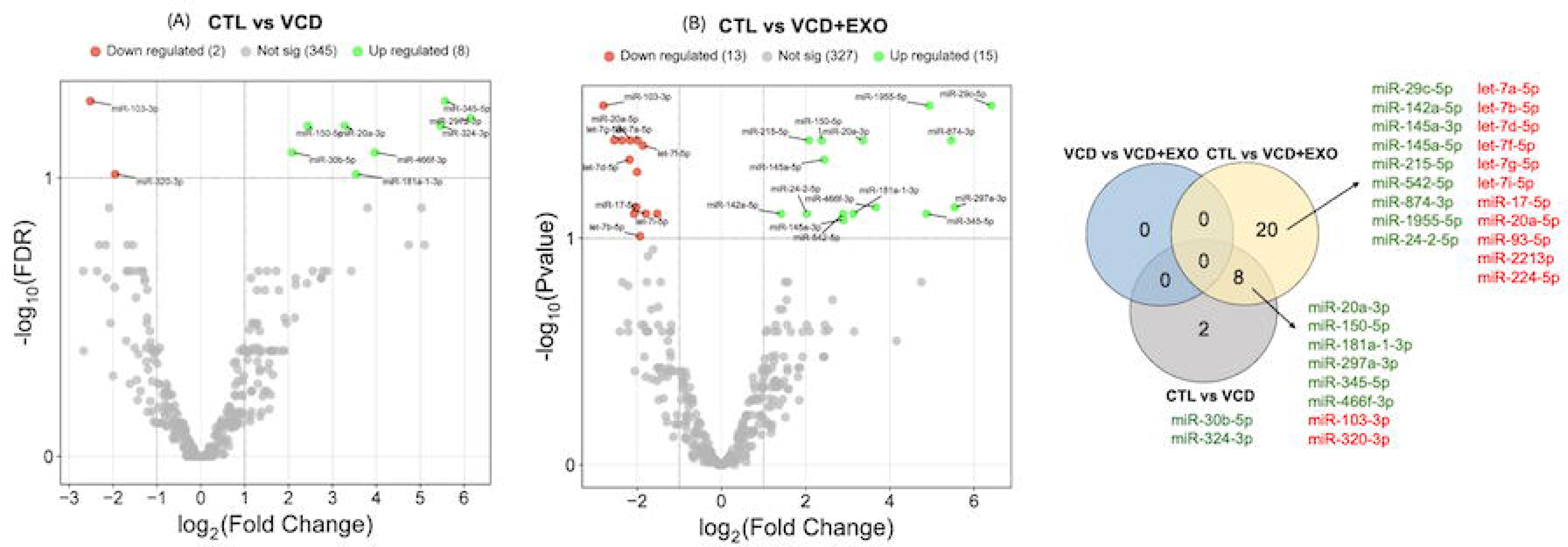
Differentially expressed serum exosomal microRNAs (miRNAs). Volcano plot with miRNAs that are significantly different colored in red for downregulated and green for upregulated for (A) CTLvsVCD and (B) CTLvsVCD+EXO. Venn diagram showing significantly regulated miRNAs among control (CTL), estropause (VCD), and estropause exosomes (VCD+EXO) mice. For each group, upregulated miRNAs (indicated in green) and downregulated miRNAs (indicated in red) are represented in the diagram.

### Liver transcriptome

We performed mRNA sequencing to compare gene expression patterns in the liver of CTL, VCD and VCD+EXO mice. We identified 13,832 genes expressed in liver. The PCA (**Fig. 7A**) and heatmap (**Fig. 7B**) analysis revealed a clear distinct profile among the three groups. In the heatmap analysis it is possible to observe that VCD+EXO and CTL groups had a more similar profile than the VCD group. This suggests that exosome treatment influence liver gene expression profile of estropause mice to be more similar to CTL cyclic mice. The number of genes regulated between groups is shown in the Venn Diagram **(Fig. 8A)** and Volcano plots **(8C-E).** We observed significant regulation of liver transcriptome induced by exosome treatment. Interestingly, seven genes were down-regulated in VCD (compared to CTL) and up-regulated in VCD+EXO (compared to VCD) mice (*Meig1, Spata4, Pde1b, Tnp1, Tnp2, Clec2m,* and *Prm2*). Conversely, two genes were up-regulated by VCD and down-regulated in VCD+EXO mice (*Lcn2* and *Mup18*). These indicate that exosome treatment could partially reverse changes induced by estropause in mice. The complete list of detected genes is shown in **Suppl. Table 3**. KEGG pathway analysis revealed that exosome treatment resulted in regulation of a significant number of pathways compared to estropause mice **(Figure 9)**. We observed that eight pathways that were up-regulated in estropause mice, were down-regulated by exosome treatment: Insulin signaling pathway, Ascorbate and aldarate metabolism, Ubiquitin mediated proteolysis, N-Glycan biosynthesis, Aminoacyl-tRNA biosynthesis, Endocytosis, Porphyrin and chlorophyll metabolism and Protein processing in endoplasmic reticulum. The Ribosome pathway was down-regulated in estropause mice and up-regulated by exosome treatment. This indicates that changes in liver transcriptome induced by estropause can be partially reversed by exosome treatment.

**Figure 7.**
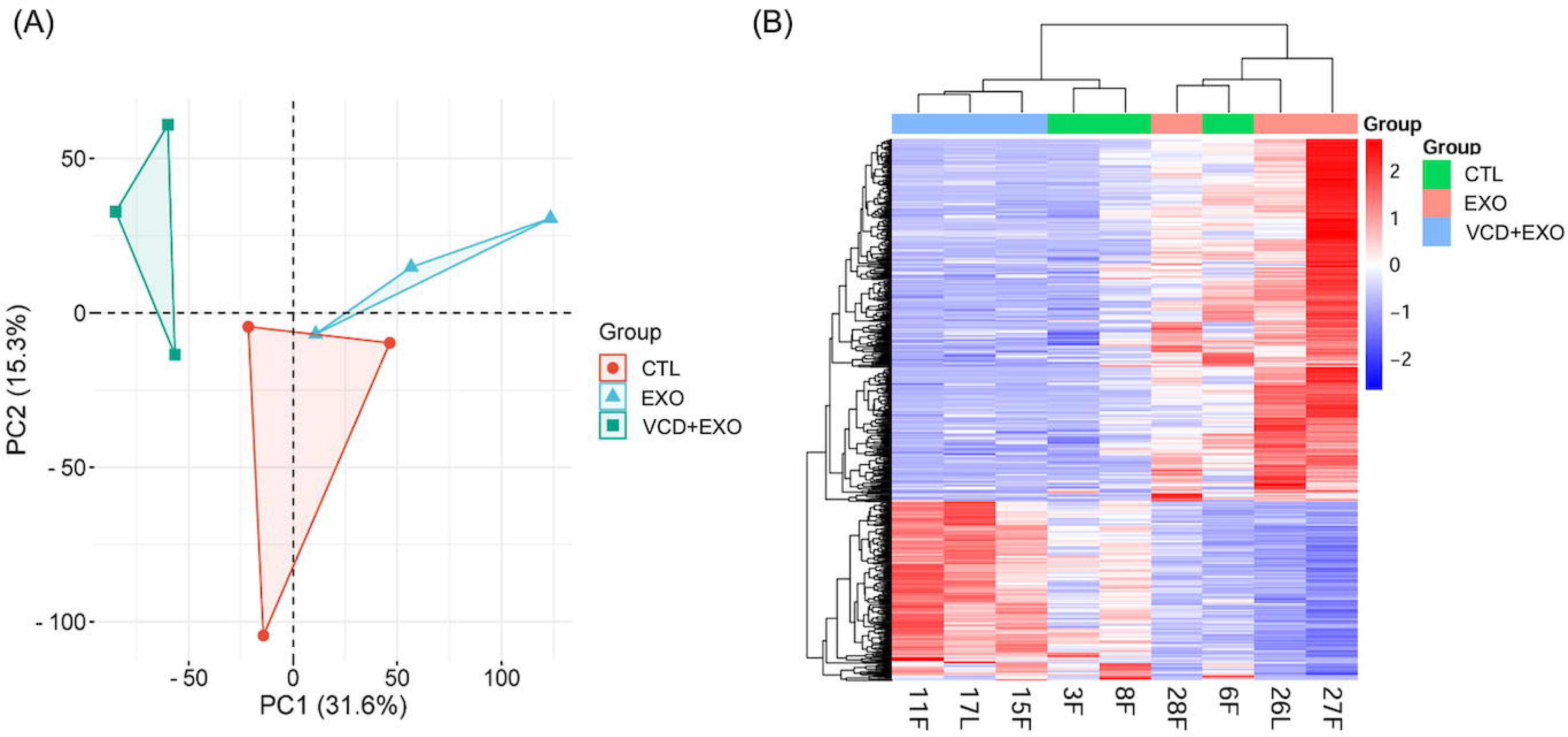
Liver transcriptome analysis. (A) Principal component analysis of liver transcriptome for control (CTL), estropause (VCD)and estropause exosomes (VCD+EXO) mice; (B) Unsupervised hierarchical clustering ordered by the adjusted level of mRNA expression for all expressed mRNA from liver tissue.

**Figure 8.**
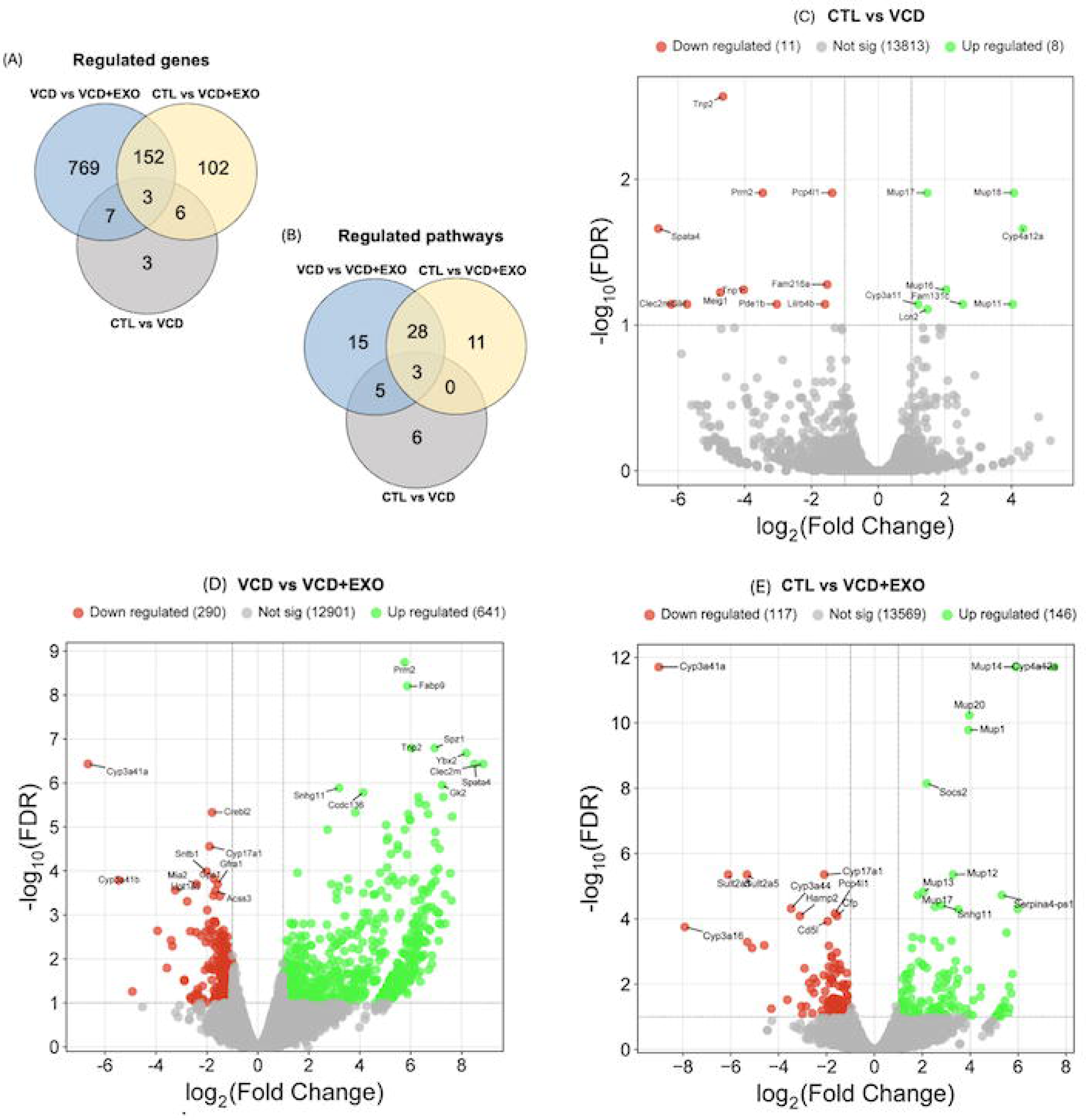
Differentially expressed liver genes. Venn diagram indicating significantly regulated genes (A) and pathways (B) from liver tissue between control (CTL), estropause (VCD) and estropause exossomes (VCD+EXO) mice. Volcano plot with significantly regulated genes colored in red indicating downregulated and green indicating upregulated for (C) CTLvsVCD; (D) VCDvsVCD+EXO and (E) CTLvsVCD+EXO.

**Figure 9.**
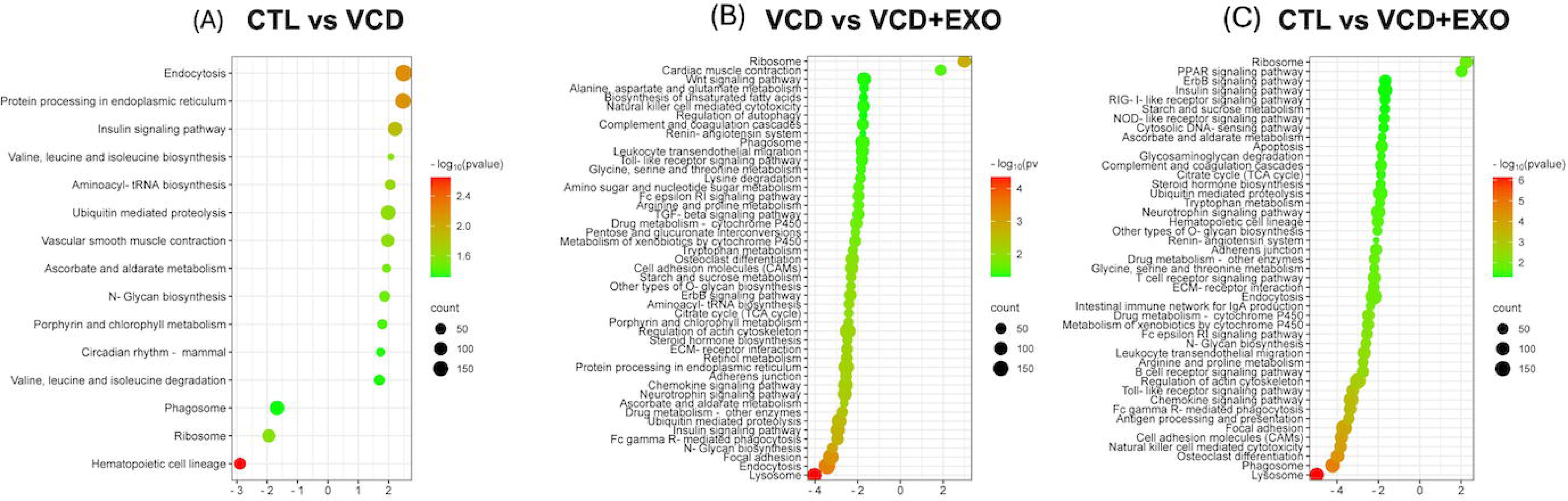
Liver regulated KEGG pathway analysis. Significant regulated pathways in liver tissue among control (CTL), estropause (VCD) and estropause exossomes (EXO) mice. (A) CTLvsVCD (B) VCDvsVCD+EXO and (C) CTLvsVCD+EXO.

### Comparative serum miRNAs and liver transcriptome analysis

The genes *Cyp3a11* and *Cyp4a12a* were up-regulated in the liver of VCD and VCD+EXO compared to CTL mice. Both genes are predicted targets of miR-103-3p, which was down-regulated in serum exosomes from both VCD and VCD+EXO mice compared to CTL. Additionally, miR-224-5p and miR-20a-5p were down-regulated in VCD+EXO compared to VCD mice. *Cyp3a11* and *Cyp4a12a* are also predicted targets of these two miRNAs.

Interestingly, when observing miRNA regulated in serum exosomes from VCD compared to CTL mice, two up-regulated miRNAs are not up-regulated in the VCD+EXO compared to CTL group (miR-324-3p, miR-30b-5p). These could suggest beneficial effects of exosome treatment in modulation of these miRNAs levels. Interestingly, the predicted target genes of these miRNAs were regulated in liver of VCD+EXO compared to VCD mice. This reinforces a positive role of exosomes in the regulation of liver transcriptome in estropausal mice.

## Discussion

We observed that estropause induction in young mice leads to a small increase in body and fat mass and minor changes in liver transcriptome. This is interesting and suggests that in the absence of aging, cessation of ovarian function alone leads to few metabolic adaptations in females. We previously also observed a small increase in body weight and no changes in glucose metabolism or oxidative status in liver of young VCD treated mice [28,32]. Others also showed no changes in body mass and only small changes in liver transcriptome after estropause induction by VCD [33]. We detected *Mup11*, *Mup16*, *Mup17* and *Mup18* as up-regulated genes in liver by VCD, similar to a previous study, which observed *Mup15* and *Mup17* as up-regulated by VCD [33]. Others also detected *Mup3*, *Mup7*, *Mup11*, *Mup12*, *Mup21* as up-regulated in liver of ovariectomized mice [51]. The major urinary protein (MUP) family members are produced in the liver, cand bind volatile compounds and are secreted in the urine. Some circulating MUPs have a role in glucose and lipid metabolism [52], pointing to a possible role of these proteins in estropause regulated metabolic functions. Additionally, females produce pheromone-carrying MUPs in a cycling manner, with the highest levels during proestrus and estrous [53]. Therefore, the up-regulation of these genes in VCD mice may be related to compensatory changes due cessation of ovarian steroid production. Genes involved in drug metabolism and clearance, such as cytochrome P450 (CYPs), exhibit sex-specific expression in rodents [54-56]. We observed that *Cyp4a12a* was up-regulated in estropause females. *Cyp4a12* is highly expressed in male liver at the mRNA and protein level when compared to females in several mouse strains [57]. Interestingly, *Cyp4a12a* was almost undetectable in cyclic females in our study, suggesting that this male-specific pattern is developed upon ovarian failure. Others detected strong up-regulation of *Cyp4a12a* in ovariectomized females to be mediated by ERα [51], suggesting a role of estrogens in liver masculinization after estropause. *Cyp4a12a* participates in the metabolism of arachdonic acid and regulates its circulating levels [58]. Although changes in hepatic transcriptome resulting from estropause induction were small, they indicate that the loss of ovarian function leads to masculinization of the liver transcriptome. This is interesting as male mice are more prone to develop liver disease compared to female mice [59]. Similarly, VCD treated female mice develop a more severe hepatic damage in response to a high-fat high-sucrose diet [33].

Exosomes play a crucial role in liver physiology and pathophysiology. Multiple effects of exosomes on cell-to-cell communication in the liver were demonstrated, leading to regulation of inflammation, angiogenesis, proliferation, and tissue remodeling [60-63]. We observed that exosome treatment led to significant changes in liver transcriptome, with over 900 genes regulated in comparison to estropause placebo mice. Among these, we highlight genes regulated in the opposite direction, i.e. down-regulated in estropause (VCD) compared to cyclic mice (CTL), but up-regulated in exosome treated estropause (VCD+EXO) compared to placebo treated estropause mice (VCD). These changes suggest that exosome treatment was able to partially rescue the liver transcriptomic profile of cyclic mice. In this sense, we observed only 260 genes regulated in VCD+EXO compared to CTL mice. Interestingly, we detected that *Mup18* as up-regulated in liver by estropause and down-regulated by exosome treatment, suggesting a role of exosome treatment in partially preventing liver masculinization after estropause. However, the expression of *Cyp4a12a* was up-regulated in VCD+EXO compared to VCD mice, indicating that some changes induced by exosomes can be negative. The genes *Meig1, Spata4, Tnp1, Tnp2*, and *Prm2* were down-regulated by estropause and up-regulated after exosome treatment. These were not previously observed as regulated in the VCD estropause [33] or ovariectomized models [51]. However, *Spata4* has been shown to improve aging-induced metabolic dysfunction by promoting preadipocyte differentiation and adipose tissue expansion [64]. This suggests that exosomes may regulate lipid accumulation in a tissue-specific manner through *Spata4* modulation. *Tnp1*, *Tnp2* and *Prm2* are highly expressed in testes and involved in the substitution of histone for protamines during spermiogenesis [65]. Some studies showed a role of protamine supplementation in regulation of liver metabolism, reducing serum cholesterol and body mass gain in rodents [66]. However, more studies are needed to confirm the role of these genes in liver metabolism and can be related to male specific patterns of gene expression as observed for other genes.

During menopause, hormonal changes impact metabolic pathways in the liver, contributing to changes in cholesterol, HDL and triglycerides levels and susceptibility to cardiovascular diseases [67]. Looking at liver regulated pathways, we observed that the insulin signaling pathway was up-regulated in estropause mice and down-regulated by exosome treatment. The insulin signaling pathway is essential for modulation of hepatic gluconeogenesis and lipogenesis [68]. Regulation of insulin signaling pathway was observed in the liver of ovariectomized mice that developed steatosis compared to cyclic mice [69]. This suggests that liver damage after estropause may be mediated by insulin signaling. The ubiquitin-mediated proteolysis pathway is essential for the degradation of damaged proteins, directly affecting liver function [70]. We observed the ubiquitin pathway to be upregulated in VCD compared to CTL mice, suggesting an important role in the stress response.

Ubiquitination is up-regulated in mice in response to CCl_4_ induced liver fibrosis in mice [71] and the ubiquitinated intermediate filament body inclusions are considered biomarkers of liver disease [72]. Therefore, down-regulation of the ubiquitin pathway by exosomes can indicate a protective effect in liver damage. The N-glycan pathway was also upregulated in VCD compared to CTL mice and reverted by exosome treatment. The induction of liver damage in mice by a western diet resulted in increased expression of N-glycan products in the liver tissue [73]. The sites of N-glycan release correlate with regions of increased steatosis [73]. Therefore, the regulation of this pathway in estropause mice further suggests the increased susceptibility to liver damage that was prevented by exosomes. We also observed that protein processing in the endoplasmic reticulum (ER) pathway was up-regulated by estropause and reduced by exosomes treatment. This pathway encompasses several genes involved in protein ubiquitination and N-glycosylation, therefore, in line with previous discussion. The ribosome pathway was down-regulated in estropause mice and up-regulated by exosome treatment. Interestingly, this is also directly linked to the pathogenesis of liver disease. Knockout of genes that impair ribosome biogenesis in liver of adult mice can induce hepatocyte cell cycle arrest [74] and increased liver damage with age [75], suggesting a negative effect in liver regeneration. Overall, the changes observed in liver pathways induced by estropause result in detrimental effects, which were effectively reversed by injection of exosomes.

We observed exosomes injected i.p in blood smears 24 hs after injection, indicating their delivery into circulation. Few data on circulating miRNAs and exo-miRs are available in estropausal mice and even menopausal women. We observed that eight miRNAs were regulated in estropausal mice independent of exosome treatment. Among these, miR-320-3p was also down-regulated in estropause females from our previous study [28]. Although circulating miRNAs have been extensively examined as cancer biomarkers [76], the role of miRNAs in aging, particularly in menopause, has only recently begun to be explored. Comparing the miRNAs regulated in serum of pre-and postmenopausal women [29] with the ones from our study, we found none to be commonly regulated. This may suggest a different profile between species or the confounding effects of age when evaluating pre and post-menopausal women. Here, we provide evidence that circulating exo-miRs in serum are differentially expressed between mice in estropause compared to cyclic mice of the same age. Overall, 10 miRNAs were differentially expressed in estropause compared to cyclic mouse. In this way, we attempted to associate how changes in liver gene expression profiles may be linked to circulating exosomal miRNAs. Among predicted target pathways of miRNAs regulated by estropause we observed insulin signaling pathway and valine, leucine and isoleucine degradation, which were observed as regulated pathways in the liver transcriptome in the same direction. These suggest serum miRNA changes can regulate liver transcriptome and that further functional studies are needed. Importantly, miR-103-3p was down-regulated in serum exosomes of estropause mice and has *Cyp4a12a* as a predicted target gene. We observed a strong up-regulation of *Cyp4a12a* in liver in estropause females. As previously discussed, *Cyp4a12a* is a male-specific liver gene, and our data suggests that miR-103 may play a role in its regulation after estropause. Others also reported reduced miR-103 expression in serum after OVX [77], indicating that estradiol may promote this sex bias. Functional studies are needed to confirm the role of miR-103 in the regulation of *Cyp4a12a*.

We observed that miR-30b-5p and miR-324-3p miRNAs were exclusively up-regulated in VCD compared to CTL mice and 20 miRNAs were exclusively regulated in VCD+EXO compared to CTL mice. Previous studies observed that miR-30b is more expressed in the blood of men than women [78]. This suggest a possible masculinization of the serum miRNA profile that is partially reversed by exosome treatment. miR-324-3p was found to be upregulated in postmenopausal women and associated with bone mineral density [79]. Although no difference was detected in serum miRNAs between placebo and exosome treated estropause mice, the miRNAs exclusively regulated in VCD+EXO mice suggest a small residual effect of exosome treatment 48 h after injection. miR-345-5p was up-regulated in VCD and in VCD+EXO treated mice. miR-345-5p is involved in adipocyte differentiation by suppressing the expression of VEGF-B [80]. VEGF signaling pathway was observed as exclusively regulated by exosomes in liver in our study. VEGF-B induces the expression of fatty acid transport protein 3 (*FATP3*) and *FATP4*, which are regulators of fatty acid metabolism in liver [81]. miR-181a was up-regulated in VCD and VCD+EXO mice and regulates E2 signaling and is implicated in hormone-dependent cancers [82]. In some conditions E2 can increase miR-181a-5p expression [82]. Interestingly, the let7 family was down-regulated exclusively in exosome treated mice (7a, &b, 7d, 7f, 7g and 7i). Lower let-7 expression is associated to poor cancer prognosis [83] and to liver damage [84]. Our findings therefore demonstrate that cyclicity is a critical regulator of exosomal miRNA levels and can have implication in development of diseases.

In conclusion, we observed that estropause in young mice induced few metabolic changes, with a slightly increased body and fat mass. This allowed us to isolate the effect of ovarian failure from aging and measure its direct effects on the exosome profile and liver transcriptome. Changes in liver transcriptome were minor in estropausal mice but indicate a masculinization of the gene expression pattern. Exosome treatment led to drastic changes in liver transcriptome, partially reversing changes induced by estropause in pathways involved in insulin signaling and N-Glycan biosynthesis. Our findings indicate that exosomes can modulate metabolic health in estropausal females. Further functional studies are still needed to develop exosome-based therapies.

## Data availability

The data is available from the corresponding author upon request. Raw miRNA and mRNA sequencing data is available from NCBI SRA PRJNA1171446.

## Author contribution

**Bianka M Zanini,** Conceptualization; Data curation; Formal analysis, Writing - original draft **Bianca M Ávila,** Investigation, Visualization, Writing - review & editing **Jéssica D Hense,** Investigation, Visualization, Writing - review & editing **Driele N Garcia,** Investigation, Visualization, Writing - review & editing **Sarah Ashiqueali,** Investigation, Visualization, Writing - review & editing **Pâmela I. C. Alves,** Investigation, Visualization, Writing - review & editing **Thais L Oliveira,** Investigation, Visualization, Writing - review & editing **Tiago V Collares,** Investigation, Visualization, Writing - review & editing **Miguel A. Brieño-Enríquez,** Conceptualization, Writing - review & editing **Jeffrey B Mason,** Conceptualization, Writing - review & editing **Michal M Masternak,** Conceptualization, Funding Acquisition, Writing - review & editing **Augusto Schneider** Conceptualization, Project administration, Funding Acquisition, Investigation, Visualization, Writing - review & editing

